# Adult lifespan trajectories of neuromagnetic signals and interrelations with cortical thickness

**DOI:** 10.1101/2022.10.23.513274

**Authors:** Christina Stier, Christoph Braun, Niels K. Focke

**Affiliations:** Clinic of Neurology, University Medical Center Göttingen, Göttingen, Germany; Institute for Biomagnetism and Biosignalanalysis, University of Münster, Münster, Germany; MEG-Center, Hertie Institute for Clinical Brain Research, University of Tübingen, Tübingen, Germany; CIMeC, Center for Mind/Brain Sciences, University of Trento, Rovereto, Italy

**Keywords:** magnetoencephalography, resting-state, functional connectivity, structure-function coupling

## Abstract

Oscillatory power and phase synchronization map neuronal dynamics and are commonly studied to differentiate the healthy and diseased brain. Yet, little is known about the course and spatial variability of these features from early adulthood into old age. Leveraging magnetoencephalography (MEG) resting-state data in a cross-sectional adult sample (n = 350), we probed lifespan differences (18-88 years) in connectivity and power and interaction effects with sex. Building upon recent attempts to link brain structure and function, we tested the spatial correspondence between age effects on cortical thickness and those on functional networks. We further probed a direct structure-function relationship at the level of the study sample. We found MEG frequency-specific patterns with age and divergence between sexes in low frequencies. Connectivity and power exhibited distinct linear trajectories or turning points at midlife that might reflect different physiological processes. In the delta and beta bands, these age effects corresponded to those on cortical thickness, pointing to co-variation between the modalities across the lifespan. Structure-function coupling was frequency-dependent and observed in unimodal or multimodal regions. Altogether, we provide a comprehensive overview of the topographic functional profile of adulthood that can form a basis for neurocognitive and clinical investigations. This study further sheds new light on how the brain’s structural architecture relates to fast oscillatory activity.

## 1. Introduction

Brain development and aging are subject to highly complex processes that are shaped by genetic and environmental influences and are critical factors in health and disease. The onset of various disorders often coincides with specific age windows, indicating alterations in developmental or aging pathways and genetic factors (e.g., Ellis et al. (2019); Skene et al. (2017)). For example, genetic epilepsies and psychiatric disorders such as schizophrenia occur in childhood or adolescence, while neurodegenerative diseases like Alzheimer’s disease typically emerge in the last decades of life. Efforts to quantify biological aging and normative modeling using neuroimaging for the assessment and prediction of individual health risks has gained momentum in the last decade (Cole et al., 2019), but have been primarily focused on neuroanatomical data (Bethlehem et al., 2022; Kaufmann et al., 2019). However, many pathological states such as schizophrenia or epilepsy are characterized by aberrant fast neuronal activity and synchronization that can be measured using MEG or EEG (Elshahabi et al., 2015; Hegner et al., 2018; Hirvonen et al., 2017; Kurimoto et al., 2008; Stam et al., 2002; Stier et al., 2021). Thus, understanding normative brain maturation and aging based on neuronal activity is key to estimating pathological trajectories.

Research on age effects on MEG or EEG signal power was undertaken early on (Duffy et al., 1984), providing evidence for a shift from lower to higher frequency bands (Coquelet et al., 2020; Gómez et al., 2013; Hunt et al., 2019; Marek et al., 2018; Miskovic et al., 2015; Whitaker et al., 2016), that is, a decrease in low-frequency power with age and an increase in higher frequencies. These seminal studies, however, have focused only on specific age decades (Coquelet et al., 2020; Hunt et al., 2019; Marek et al., 2018; Schäfer et al., 2014) or were done at the sensor level (Gómez et al., 2013; Sahoo et al., 2020), limiting the spatial specificity and possibility of studying anatomically distributed networks across the lifespan. Similarly, although measures of phase-synchronization have been popular in the search for biomarkers and description of cognitive processes (Sadaghiani et al., 2022), very few studies have investigated phase-based neuronal organization across a broad age range (Hunt et al., 2016). Others used sensor signal coherence as their connectivity measure (Sahoo et al., 2020), which is likely to be influenced by electromagnetic field spread.

Hence, the primary goal of this study was to investigate regional variability of oscillatory activity during the resting-state and across the entire adulthood using sophisticated source reconstruction methods to study neural generators. We hereby focused on two characteristics of neural signals such as signal power and phase-based connectivity using the imaginary part of coherency, a measure with less sensitivity to volume conduction (Nolte et al., 2004). Previous MEG studies have reported linear and non-linear relationships between age and neural signals at different frequencies (Gómez et al., 2013; Hunt et al., 2019; Rempe et al., 2023). Thus, it can be expected that neuronal activation patterns change in diverse brain regions at different time points in life. Cortical regions undergo different maturation and degeneration periods and show particular age windows for vulnerability to disruptions throughout life (Sydnor et al., 2021). For example, fMRI research shows that a gradual consolidation of brain networks is attributed to the early development stages, with association hubs becoming more strongly connected until adulthood (Oldham et al., 2022). With old age, functional connectivity within brain networks as studied using fMRI, appears to decrease, particularly in the default mode network (DMN), and relates to cognitive decline (Damoiseaux, 2017; Ferreira and Busatto, 2013). Importantly, cerebrovascular function changes with age and can substantially confound fMRI connectivity measures if not corrected appropriately (Tsvetanov et al., 2015). MEG, on the other hand, is less sensitive to vascular confounds (Tsvetanov et al., 2015), measures neuronal activity more directly than fMRI and can capture fast neuronal dynamics.

A second goal of this study was to examine potential differences between sexes in the lifespan trajectories for the MEG measures, which can be expected based on the following evidence: Males and females show divergent biological variations, for instance, in brain volume (Ritchie et al., 2018; Ruigrok et al., 2014) and fMRI network connectivity (Satterthwaite et al., 2015). Some studies report sex differences in MEG (Azanova et al., 2021; Fung et al., 2021; Hoshi and Shigihara, 2020; Taylor et al., 2020) or EEG features (Brenner et al., 1995; Clarke et al., 2001; Davidson et al., 1976; Kober and Neuper, 2011; Smit et al., 2008; Thordstein et al., 2006) during or in absence of a task in various age ranges and frequencies. However, research efforts in this direction for the resting-state across the lifespan are generally limited (Rempe et al., 2023). Providing normative maps of neuronal signatures with age and split for sex may be particularly crucial for studies in sex and gender-based medicine or cognitive phenomena.

Finally, we aimed at quantifying how the lifetime changes in neuronal patterns relate to those in brain structure. A growing body of literature has established correlations between co-activation patterns and structural connectivity using MR imaging (Suárez et al., 2020), but less often in context with aging (Park et al., 2022). Evidence from fMRI studies suggests that structure-function coupling in general (Medaglia et al., 2018) and its development during youth relates to functional and cognitive flexibility (Baum et al., 2020). As such, investigating which brain features are similarly age-dependent contribute to understanding perturbations in cognitive tasks (Suárez et al., 2020) and deviations in pathological states that often affect both structural and functional circuits (e.g., Stier et al. (2022)). However, the relation of haemodynamic connectivity during rest to MEG networks is complex and does not follow simple linear dynamics for single frequency bands (Shafiei et al., 2022; Tewarie et al., 2016). At the same time, a close relationship between beta oscillations and brain structural properties is likely, since MEG signals in this frequency range contributed most to fMRI networks (Shafiei et al., 2022; Tewarie et al., 2016). In our study, we sought to bring together lifespan patterns for fast neuronal activity measured using MEG and MRI-derived cortical thickness as index of the brain’s macrostructural organization. Gradual cortical thinning from childhood up into old age has been consistently reported, with a few exceptions for different age effects in the anterior cingulate cortex, entorhinal, and temporopolar cortices (Frangou et al., 2022). Overall, this study set out to track the brain’s functional profile in 18- to 88-year-old individuals using source-reconstructed popular MEG measures (Sadaghiani et al., 2022) and to investigate differences among sexes and intersections with cortical thickness. We hypothesized linear and non-linear associations between age and MEG power and connectivity, respectively, resolved for six frequency bands with spatially distinct regional patterns. To exclude the possibility that our age-related MEG and cortical thickness findings are confounded by intracranial volume variability (Leonard et al., 2008), we incorporated it as a factor into our statistical analyses together with sex.

## 2. Materials and Methods

### 2.1. Data and participants

In our study, we used cross-sectional open-access data provided by the Cambridge Centre for Aging and Neuroscience (Cam-CAN) data repository. The Cam-CAN project encompassed several phases, including cognitive assessments, interviews and health and lifestyle questionnaires, and structural and functional brain examinations (Shafto et al., 2014). 650 resting-state MEG data sets and T1-weighted MR-images were available from phase two of the study (Taylor et al., 2017), a subset of which we initially analyzed (*n* = 450). After excluding data with motion artifacts, noise, sleep, or failed Freesurfer reconstruction (*n*_excluded_ = 72), 350 cleaned MEG and MRI data were randomly selected from the remaining data sets for further analysis in a balanced design. In total, we report on seven age groups, each with 50 individuals aged 18 to 88 years, divided into ten-year increments and balanced by sex (see demographic data in **Table 1**). All included individuals were cognitively normal (Mini Mental State Examination score < 24; Folstein et al. (1975)) and free of neurological or psychiatric conditions (e.g. dementia, epilepsy, head injury with severe sequelae, bipolar disorder, schizophrenia) and substance abuse history. Individuals with communication problems (hearing, speech, or visual impairment), limited mobility, or MRI/MEG contraindications were excluded. For details on the study protocol and datasets see Shafto et al. (2014). The study was conducted in accordance with the Declaration of Helsinki (World Medical Association, 2013) and approved by the local ethics committee, Cambridgeshire 2 Research Ethics Committee. Data collection and sharing for this project was provided by the Cam-CAN (https://camcan-archive.mrc-cbu.cam.ac.uk/dataaccess/).

### 2.2. MRI acquisition

Anatomical data were acquired using a 3T Siemens TIM Trio scanner with a 32-channel head coil. T1-weighted images were derived from 3D MPRAGE sequences with TR=2250ms, TE=2.99ms, TI=900ms; FA=9 deg; FOV=256×240×192mm; 1mm isotropic; GRAPPA=2; TA=4mins 32s), and T2-weighted images from 3D SPACE sequences with TR=2800ms, TE=408ms, TI=900ms; FOV=256×256×192mm; 1mm isotropic; GRAPPA=2; TA=4mins 30s).

### 2.3. MEG acquisition

Resting-state data were recorded using a 306-channel VectorView MEG system (Elekta Neuromag, Helsinki) with 102 magnetometers and 204 planar gradiometers (sampling at 1 kHz with a 0.03 Hz high pass filter). Individuals were assessed in a seated position in a magnetically shielded room at a single site (MRC Cognition and Brain Science Unit, University of Cambridge, UK) for 8 min and 40 s. At the same time, four coils continuously measured the head position within the MEG helmet. Additionally, electrocardiogram (ECG) and electrooculogram (EOG, horizontal and vertical) were recorded to track cardiac signals and eye-movements. Individuals were instructed to keep their eyes closed and sit still.

### 2.4. MRI processing and individual head models

For mapping MEG sensor level data onto individual cortical surfaces, anatomical information was derived from T1- and T2-weighted images and reconstructed using FreeSurfer 6.0.0 (https://surfer.nmr.mgh.harvard.edu/). We applied surface-based mapping (SUMA; Saad and Reynolds (2012)), which resampled the cortical surfaces to 1,002 vertices per hemisphere (ld factor = 10), based on the ‘fsaverage’ template mesh provided by FreeSurfer and SUMA. The individual meshes were realigned to the Neuromag sensor space based on anatomical landmark coordinates provided by Cam-CAN. We used the “single shell” method implemented in Fieldtrip to compute the leadfields and individual head models for MEG source projection.

### 2.5. MEG processing

Preprocessed MEG data was available through the Cam-CAN repository. For each dataset Elekta Neuromag Maxfilter 2.2 was applied using temporal signal space separation (10s window, 0.98 correlation limit) to remove external interference and artefacts, line noise (50 Hz and its harmonics), and to correct for bad channels and head movements. Using Fieldtrip (Oostenveld et al., 2011), we resampled the data to 300 Hz, initially high-pass filtered at 1 Hz (first order Butterworth), and segmented the data into trials of 10 s length. Trials containing artifacts were removed following an automatic approach for both MEG channel types separately (see https://www.fieldtriptoolbox.org/tutorial/automatic_artifact_rejection/ for further details). In brief, the data was bandpass filtered at 110 to 140 Hz (9th order Butterworth) for optimal detection of muscle artifacts and z-transformed for each channel and timepoint. The z-transformed values were averaged over all channels so that artifacts accumulated and could be detected in a time course representing standardized deviations from the mean of all channels. Finally, all time points that belonged to the artifact were marked using artifact padding, and data trials whose z-values were above a threshold of 14 were excluded. The remaining data were then low-pass filtered at 70 Hz (first-order Butterworth), and independent component analysis (ICA) was applied to identify ocular and cardiac artifacts. Ocular components were automatically identified based on their similarity to EOG channel signals (average coherence > 0.3 and amplitude correlation coefficient > 0.4). Cardiac components were identified when coherent with the ECG signal (average coherence > 0.3) or based on the averaged maximum peaks timelocked to the ECG (QRS complex, see https://www.fieldtriptoolbox.org/example/use_independent_component_analysis_ica_to_remove_ecg_artifacts/). For each data set, the automatic selection of the components was visually checked. In a few cases, we had to manually select the relevant ICA components because the ECG/EOG was noisy. We visually inspected all cleaned data for quality control and rated vigilance of individuals according to the criteria of the American Academy of Sleep Medicine (https://aasm.org/). Thirty trials from the cleaned data scored as “awake” (> 50% alpha activity within a trial) were randomly selected for source analysis as good reliability has been shown for the metrics of interest for five minutes of data (Marquetand et al., 2019). Only signals from magnetometers (102 channels) were used and beamforming applied (dynamic imaging of coherent sources; Gross et al. (2001)) to project sensor data to the surface points (source space) in the frequency domain. MEG power and cross-spectral densities were computed for six conventional frequency bands (delta: 2 ± 2 Hz, theta: 6 ± 2 Hz, alpha 10 ± 2 Hz, low beta 16 ± 4 Hz, high beta 25 ± 4 Hz and gamma 40 ± 8 Hz) based on fast Fourier spectral analysis using multitapers (Discrete Prolate Spheroidal Sequences tapers). Source projection was carried out using leadfields and adaptive spatial filters (regularization: lambda = 5%). The coherency coefficient was estimated between all pairs of vertices (source points, *n* = 2238) and the imaginary part was derived to account for potential field spread (Nolte et al., 2004). The absolute imaginary part of coherency was our connectivity measure of interest, reflecting phase synchrony between signals. We averaged all connections of a vertex to obtain the overall strength of a vertex, and for each individual also across all vertices to get a global connectivity and power index. To provide an overview of connectivity and power distributions across age groups, the frequency spectra of each individual in this study were calculated for 1-Hz bins and averaged for young, middle-aged, and old participants (**Suppl Figure 1**).

### 2.6. Cortical thickness estimations

At each vertex of the cortical surface derived from the Freesurfer/SUMA procedure, individual cortical thickness measures were calculated, which describe the distance between the gray-white matter and pial boundaries. Smoothing of individual thickness maps were applied using a heat kernel of size 12 mm full width at half maximum in AFNI (https://afni.nimh.nih.gov/).

### 2.7. Permutation-based analysis of lifespan and sex differences

Previous studies have reported linear and non-linear relationships between age and MEG measures and better performance of quadratic regression than linear regression (Gómez et al., 2013). We therefore examined the relationship between age and the imaging metrics (connectivity, power, or cortical thickness) by fitting linear models with the metrics as dependent variables and age, age^2^, sex, and intracranial volume as independent variables. We mean centered the individual age values before squaring them to reduce the correlation between the linear and the quadratic terms. The full model was estimated either for the global metrics or for the metrics at each vertex (surface point) in the brain and in each frequency band separately. The non-parametric statistic tool PALM (Permutation Analysis of Linear Models, (2014)) was used to generate permutations for the respective models with tail approximation for accelerated inference (500 permutations) (Winkler et al., 2016). Single *t* contrasts were computed, that is, for positive and negative linear age effects, and for convex and concave quadratic age terms. Using PALM, we also investigated sex differences in the trajectory of connectivity and power over the lifespan. For each age decade, 25 males and 25 females were included to ensure a balanced design giving 175 individuals per sex in total. Models were fitted to test whether the beta coefficients for the age and age^2^ effects on connectivity or power in each frequency band differed between males and females while correcting for the influence of total intracranial volume. For all analyses using the above described permutation approach, *p*-values were derived from the permutation distribution, at the tail of which a generalized Pareto distribution was fitted (Winkler et al., 2016), and corrected for multiple comparisons (family-wise error, FWE) at cluster level resulting from threshold-free cluster enhancement (Smith and Nichols, 2009). We set the significance level at *p* = 0.05 or equivalently -log10(*p*) ∼ 1.3. The partial correlation coefficient (*r_partial_*) was estimated as an effect size for the independent variables based on the *t*-values and degrees of freedom of the global models (Rosenthal et al., 1994). *r_partial_* indicates the degree of association between two variables at which the influence of other variables in the model has been eliminated (Bortz and Schuster, 2011): values of ± 0.1 reflect a small effect, ± 0.3 represent a moderate effect, and ± 0.5 is a large effect (Field et al., 2012).

### 2.8. Spin-tests for correspondence of age effects on brain structure and MEG markers

To statistically relate age effects on connectivity or power to those on cortical thickness, we used the unthresholded *t*-value maps derived from the main analyses on lifespan differences without the application of threshold-free cluster enhancement (Smith and Nichols, 2009). We estimated Spearman rank correlations between the *t*-maps for the linear (negative) and quadratic (inverted U-shaped) age effects on cortical thickness and the maps of the linear and quadratic age effects on connectivity and power, respectively. Statistical significance for each correlation was assessed based on null distributions generated using 10,000 random rotations of the spherical projection of the cortical surface (SUMA) (“spin-test”; Alexander-Bloch et al. (2018)). This approach ensures the spatial embeddedness of the *t*-maps, as the rotational alignment is randomized rather than assuming exchangeability of the surface points. Rotations were applied to each hemisphere separately. The significance level was set to 5%.

### 2.9. Structure-function relationships beyond age

Finally, we explored if there was an overarching link between cortical thickness and functional MEG levels when considering the entire lifespan data. Because strong structure-function correspondence can be expected in functionally relevant systems (Vázquez-Rodríguez et al., 2019), we averaged the individual cortical thickness and MEG values across surface-vertices corresponding to respective functional resting-state networks (Yeo et al., 2011). PALM was run for each of the networks on both hemispheres (500 permutations, accelerated inference (Winkler et al., 2016)) and false discovery rate (FDR) applied to correct for multiple comparisons at the 14 networks. For each network, we report the effect of cortical thickness on the MEG markers after correcting for the effects of age, age^2^, sex, and total intracranial volume. The results for the functional networks were visualized using the ‘ggsegYeo2011’ package in R Statistical Software (R Core Team, 2020).

## 3. Results

### 3.1. Lifespan differences in phase-based connectivity and effects of sex

We found that connectivity varied linearly with age in mainly posterior parts of the brain with similar topographies for the frequency bands studied, involving cuneus and inferior parietal regions (*p*_FWE_ < 0.05). The direction of the effect was positive for the theta and gamma frequency bands and, conversely, negative in the alpha to high beta bands (**Figure 1**). Despite relatively local association patterns with age, the linear effects in these frequency bands were also significant on a global level for theta, alpha, and gamma (range global *r_partial_* = 0.09-0.17, **Suppl Table 2**). Interestingly, there was a different pattern for the low and high beta bands in central and frontotemporal regions that significantly followed a quadratic function with age (**Figure 1**). That is, middle age groups exhibited the strongest beta connectivity levels in comparison with young or old age groups (vertex-wise and globally with *r_partial_* ranging from 0.18-0.21, **Suppl Table 2**). There was no evidence for main effects of sex or interaction effects for age and sex, except for delta connectivity, such that females had a steeper slope for delta connectivity than males in the cuneus and parietal regions. At a global level, the interaction was not significant for the linear age effects, but for the quadratic age term, overall pointing to differences in delta connectivity between the sexes (**Figure 3, Suppl Table 4**).

**Figure 1.**
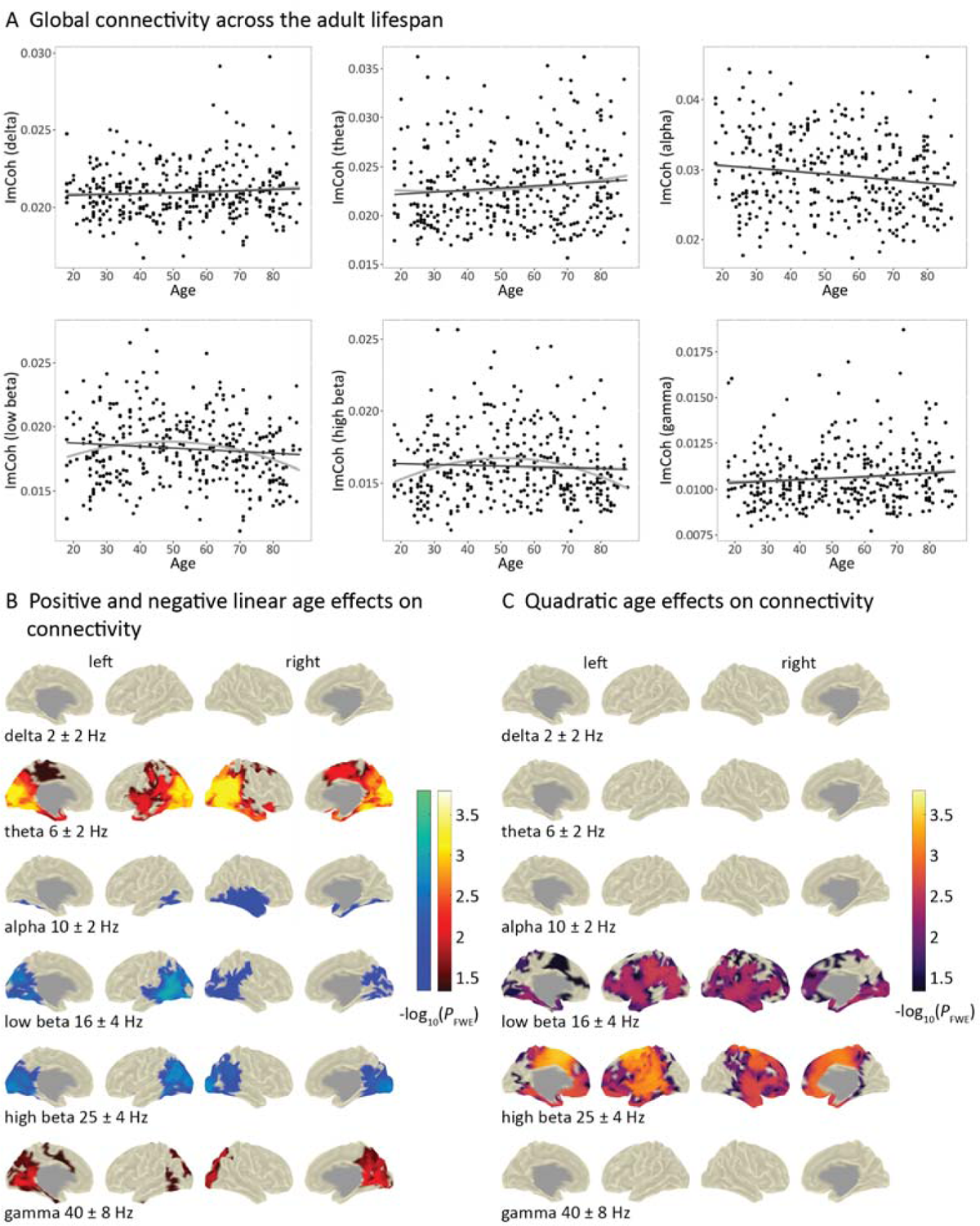
Frequency-specific associations between age and connectivity. (**A)** The plots show individual (raw) values of global connectivity (*n* = 350) across early and late adulthood for the six frequency bands investigated. The black lines represent linear relationships between age and raw global connectivity values, while the gray lines represent quadratic relationships. Statistical analyses yielded significant linear effects of age in the theta, alpha, and gamma frequency bands (range *r_partial_* = 0.09-0.17). The quadratic term in the regression model was significant for the beta1 and beta2 bands (range *r_partial_* = 0.18-0.21). See **Suppl Table 2** for statistical details. In **(B)** and **(C)**, significant effects of age on vertex connectivity are highlighted. **(B)** The blue color bar indicates significant negative effects of age, whereas the red color bar represents significant positive associations. **(C)** The purple color bar indicates significant quadratic effects of age following an inverted U-shaped pattern (concave). Results for the U-shaped term (convex) were not significant and are not displayed. The significance level was set at -log10 *p* > 1.3 (equivalent to *p* < 0.05), family-wise error corrected (FWE). We estimated linear models for each frequency band separately with connectivity as the dependent variable, and age, age^2^, sex, and intracranial volume as independent variables and performed permutation-based analysis (at a global and vertex-level). ImCoh = imaginary part of coherency. Left = left hemisphere, right = right hemisphere.

### 3.2. Lifespan differences in power and effects of sex

Age effects on power were mostly distinct from those on connectivity and tended to be focused on anterior-temporal and central regions with shifts from lower to higher frequency bands. Specifically, power in the theta to gamma bands showed a significant positive association with age, globally (range global *r_partial_* = 0.12-0.25, **Suppl Table 3**) and with most prominent effects in frontotemporal, insular, and central regions (**Figure 2**). There was also a significant negative association with global delta power (*r_partial_* = 0.11, **Suppl Table 3**), which did not survive corrections for multiple comparisons in the vertex-analysis. Furthermore, we observed quadratic age effects on power (**Figure 2**). Delta power and less strong theta power, globally showed a dip in the middle ages (range global *r_partial_* = 0.10-0.19, **Suppl Table 4**) with a strong anterior-basal emphasis, whereas the opposite direction was found for the beta and gamma bands peaking in the middle ages (range global *r_partial_*= 0.10-0.22). Of note, the quadratic age effects in high beta and gamma were prominent in the cingulate and central areas, overlapping spatially, at least in part, with the quadratic age effects on beta connectivity. Moreover, we found a significant main effect of sex in gamma power for a few clusters in the frontal cortex (data not shown). Main effects of sex for the remaining frequency bands were not significant (either vertex-wise or globally, **Suppl Table 3**). However, as with connectivity, sex-specific analyses for power revealed differing lifespan trajectories for the delta band between males and females, with a stronger linear decline for men than women. This was the case in the global analysis **(Suppl Table 4)** as well as in frontocentral regions, cingulate, and precuneus (**Figure 3**). In frontal brain areas, there was also an interaction effect between age and sex in the theta range, which was not significant at a global level (**Figure 3**; **Suppl Table 4**).

**Figure 2.**
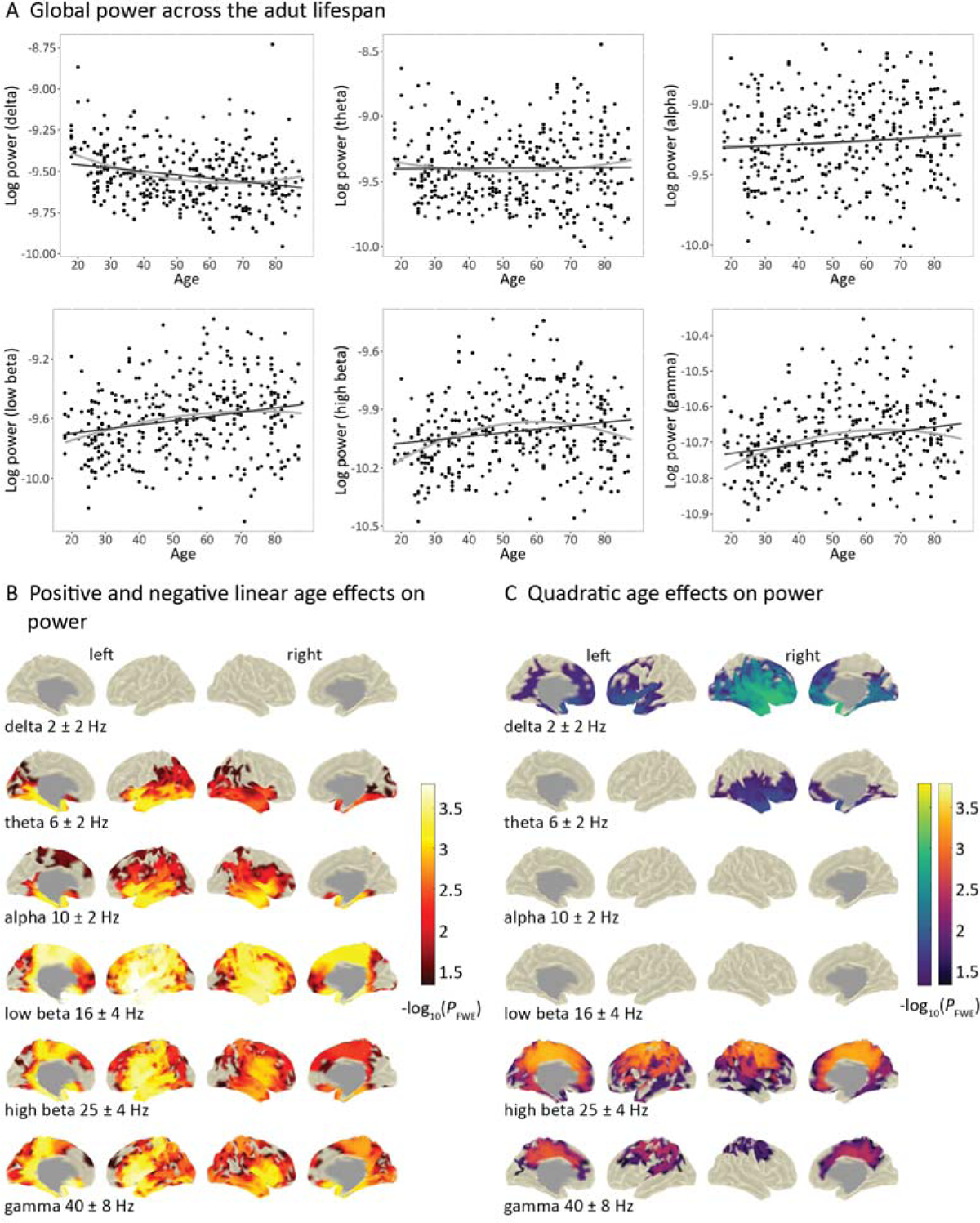
Frequency-specific associations between age and power. (**A)** The plots show individual (raw) values of global power (*n* = 350) across early and late adulthood for the six frequency bands investigated. The black lines represent linear relationships between age and raw global power, while the gray lines represent quadratic relationships. Power values (fT^2^) were log10-transformed for visualization purposes. Statistical analyses yielded significant linear effects of age on global power in all frequency bands (range *r_partial_*= 0.11-0.25). The quadratic term in the regression model was significant for the delta, theta, beta1, beta2, and gamma bands (range *r_partial_* = 0.10-0.22). See **Suppl Table 3** for statistical details. In **(B)** and **(C)**, significant effects of age on vertex-power are highlighted. **(B)** The red color bar indicates significant positive effects of age. **(C)** The green color bar depicts significant quadratic effects of age following a U-shaped pattern (convex), while the purple color bar indicates significant effects following an inverted U (concave). The significance level was set at -log10 *p* > 1.3 (equivalent to *p* < 0.05), family-wise error corrected (FWE). We estimated linear models for each frequency band separately with power as the dependent variable, and age, age^2^, sex, and intracranial volume as independent variables and performed permutation-based analysis (at a global and vertex-level). Left = left hemisphere, right = right hemisphere.

**Figure 3.**
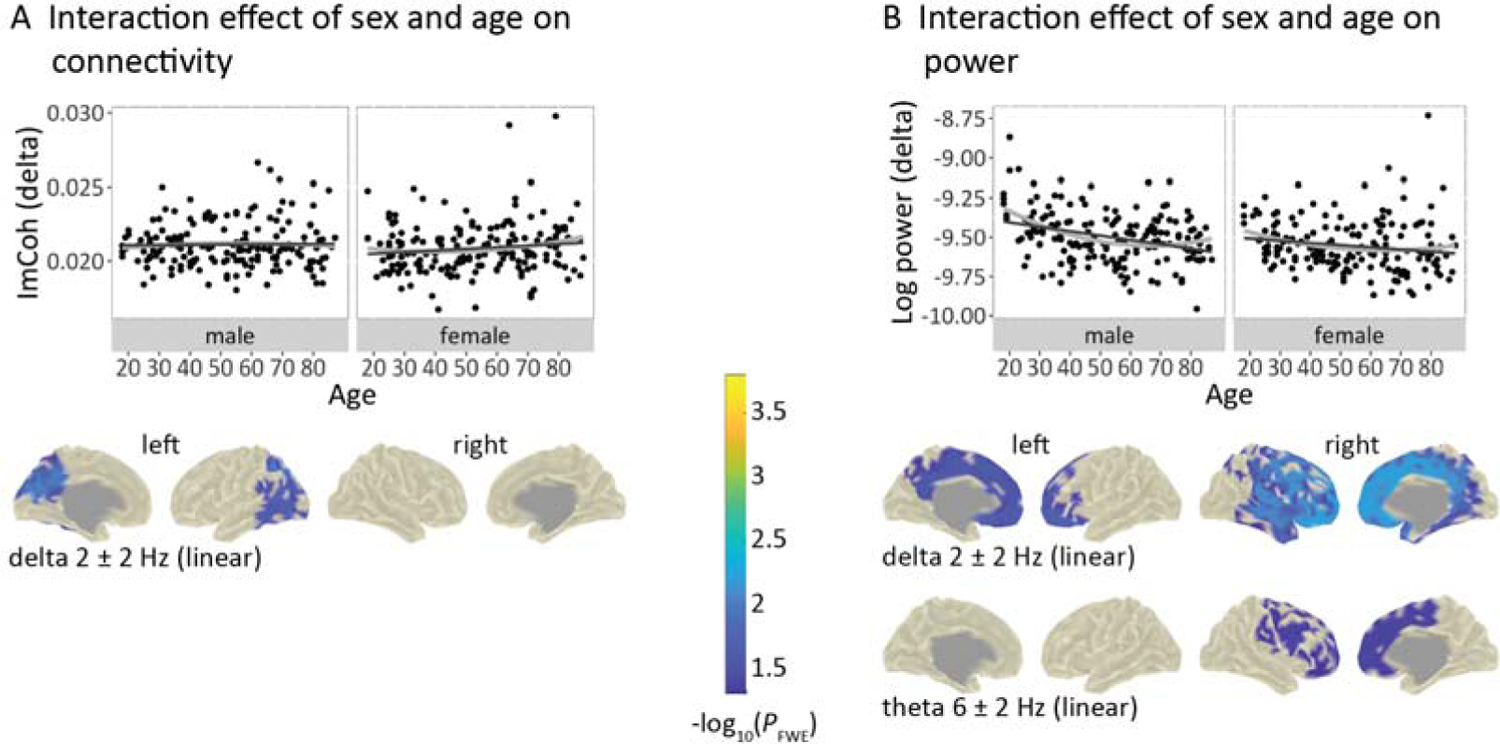
Sex-specific trajectories for MEG markers with age. **(A)** The plot shows individual global connectivity (raw values) across early and late adulthood separately for males (*n* = 175) and females (*n* = 175) in the delta band, with a significant interaction of sex and age on global connectivity (quadratic effect), such that the age effect curve was narrower for females than for males. However, at the vertex-level, the slopes for the sexes were significantly different only for the linear age effect on delta connectivity and prominent in the cuneus. **(B)** Shown are significant interactions between sex and linear age effects on global power in the delta band and on delta and theta power at the vertex level, with emphasis on frontal regions. Global power values (fT^2^) were log10-transformed for visualization purposes. The significance level was set at -log10 *p* > 1.3 (equivalent to *p* < 0.05), family-wise error corrected (FWE). We estimated linear models to test whether the age trajectories of power differed between males and females in each frequency band using permutation-based analysis. See **Suppl Table 3 and 4** for statistical details. The black lines in the global plots represent linear relationships between age and global raw MEG values, while the gray lines represent quadratic relationships. ImCoh = imaginary part of coherency. Left = left hemisphere, right = right hemisphere.

### 3.3. Assessment of structural alterations across adulthood

In accordance with previous reports (Frangou et al., 2022), a dominant pattern of a significant gradual decrease of cortical thickness with age was observed. This linear age effect was significant, globally (*p* = 0.002) and in most of the brain areas investigated (**Figure 4A**), particularly in temporal regions, supramarginal and inferior parietal regions, posterior cingulate, central, insular and caudle middle frontal regions. In a few areas, cortical thickness followed a quadratic trajectory with age, with the highest values in the middle age groups in the right temporal pole, parahippocampal and lateral occipital areas (**Figure 4A**), as previously reported (Frangou et al., 2022).

**Figure 4.**
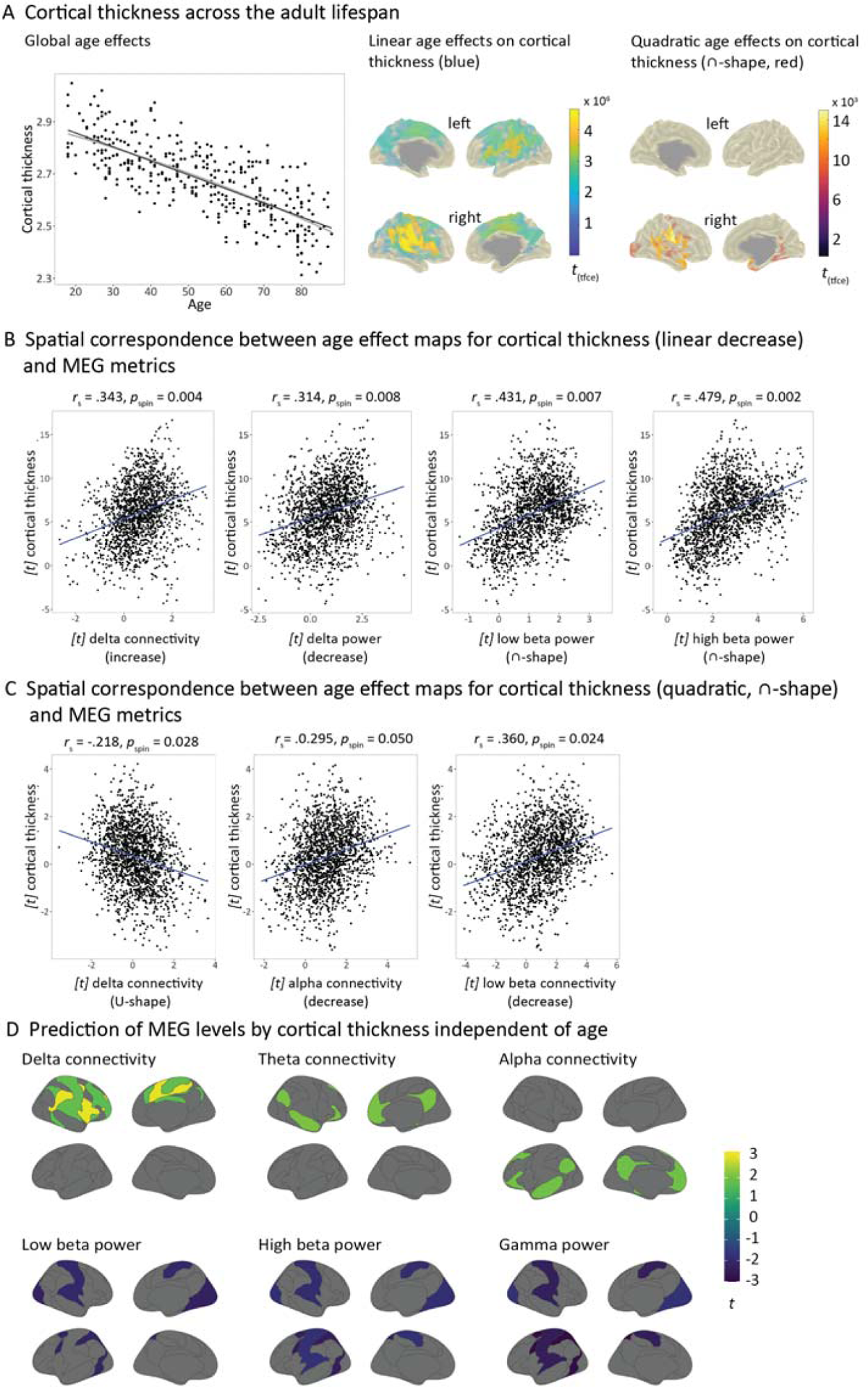
Age effects on cortical thickness and relation to MEG markers. (A) Age effects on global and vertex-wise cortical thickness across early and late adulthood. The black line represents the linear relationship between age and raw global thickness, while the gray line represents a quadratic relationship. The linear negative age effect was significant in various brain regions with an emphasis on temporal-parietal regions (blue color bar). Cortical thickness in a few clusters followed a quadratic relationship with age (red color bar, non-significant on a global level). Shown are *t*-values after threshold-free cluster enhancement (TFCE) for significant vertices. Left = left hemisphere, right = right hemisphere. (B) The gradual decrease in cortical thickness significantly correlated with age-related patterns for mostly delta (linear) and beta power (quadratic), but also for delta connectivity (linear). (C) There was also a significant spatial overlap between the quadratic age effects on cortical thickness and those on delta (quadratic), alpha, and beta connectivity (linear). Spearman rank correlation coefficients (*r*_s_) were tested for significance using a spatial permutation procedure (spin-test, 10,000 rotations). Note that the regression lines were added to the plots for visualization purposes only. (D) Across the entire adult lifespan sample, cortical thickness levels significantly aligned with connectivity levels in the delta to alpha bands. A negative effect of cortical thickness was observed for high frequency power. Only *t*-values in networks that were significant at the uncorrected level are shown. The full *t*-maps can be found in Suppl Fig. 2. Note that only the structure-function correspondence for delta connectivity in the ventral attention network and gamma power in the sensorimotor network survived corrections for the number of networks tested (FDR-correction).

### 3.4. Relating lifespan patterns for cortical thickness and MEG markers

The cross-sectional course of cortical thickness with age spatially corresponded to that of beta and delta oscillations (**Figures 4B and 4C**). Specifically, the linear decrease of cortical thickness with age was mostly correlated with age effects on MEG power, particularly in the beta (quadratic age effects low beta *r*_s_ = .431, *p*_spin_ = 0.007; high beta *r*_s_ = .479, *p*_spin_ = 0.002) and delta frequency bands (linear age effect, *r*_s_ = .314, *p*_spin_ = 0.008). This correspondence was weaker in the remaining frequency bands and did not reach significance after applying the spin test (*r*_s_ ranging from |.014-.366| for linear and quadratic age effects on power, *p*_spin_ > 0.05; **Suppl Table 5).** There was also a moderate, but significant correlation between cortical thinning and age effects on connectivity in the delta band (linear age effect, *r*_s_ = .343, *p*_spin_ = 0.004) and weaker, non-significant correlations in the remaining frequency bands (*r*_s_ ranging from |.051-.267| for linear and quadratic age effects, *p*_spin_ > 0.05; **Suppl Table 5**). Conversely, patterns of quadratic age effects on cortical thickness mainly corresponded to age effects on connectivity, namely in the low beta band (linear age effect, *r*_s_ = .360, *p*_spin_ = 0.024), delta (quadratic age effect, *r*_s_ = −.218, *p*_spin_ = 0.028), and marginally in the alpha band (linear age effect, *r*_s_ = .0.295, *p*_spin_ = 0.050). There were no other significant overlaps between the *t*-statistics for cortical thickness and connectivity in the other frequency bands (*r*_s_ ranging from |<.001-.193| for linear and quadratic age effects, *p*_spin_ > 0.05; **Suppl Table 6**) nor with power maps (*r*_s_ ranging from |.012-.289| for linear and quadratic age effects, *p*_spin_ > 0.05; **Suppl Table 6**).

### 3.5. Correspondence of oscillatory activity and brain structure beyond age

We further assessed the influence of brain structure on functional MEG markers at the level of functional resting-state networks (Yeo et al., 2011). With an exception for the delta band, there was a tendency of positive associations between cortical thickness and MEG levels in multimodal regions, and negative associations in unimodal regions (**Suppl Figure 2**). These were significant for connectivity in the lower frequency bands and power in the higher frequency bands (**Figure 4D**). Specifically, delta connectivity in the ventral attention network was predicted by cortical thickness, which was also the case for the frontoparietal and motor networks when not corrected for the number of networks tested. Similarly, there was a positive association for theta and alpha connectivity in the DMN at the uncorrected level. Conversely, cortical thickness in somatomotor and visual networks negatively predicted beta and gamma power in these regions. However, only the pattern for gamma power survived corrections for multiple comparisons.

## 4. Discussion

Using a large set of age-stratified MEG resting-state recordings, we provide crucial insights into how phase-based connectivity and power are expressed in whole-brain networks from early adulthood to old age. The markers showed distinct, frequency- and sex-specific associations with age, likely reflecting physiological processes at various adult life stages. We further provide evidence for spatial co-variation between cortical thickness and delta/beta oscillations across the lifespan, suggesting that both modalities reflect similar maturation and aging processes. Extending earlier structure-function findings based on MRI, cortical thickness predicted MEG narrowband levels across the entire study sample, positively in multimodal and negatively in unimodal networks.

### 4.1. Frequency-dependent connectivity differences with age

Our study addresses the gap that phase-based neural organization across a broad age range has been primarily reported using fMRI methods or EEG/MEG connectivity without providing spatial information on the effects (Sahoo et al., 2020). Here, phase coupling varied primarily in posterior brain regions in a linear fashion, whereas in central-temporal areas, beta band connectivity peaked at midlife. Remarkably, the linear age effects across the frequency bands tended to be local and of similar topography, with an increase in mainly the theta range and also gamma, and a decrease in alpha to high beta. Hence, connectivity alterations in these frequency bands possibly reflect different but related functional processes in posterior brain regions. The cross-sectional nature of our study and the lack of behavioral correlates only allow speculations about the underpinnings of these effects, but an attempt will be made here. Theta coupling appears to change along an anterior-posterior gradient with age, with frontal decreases during adolescence, which was related to cognitive control (Marek et al., 2018), and increases in posterior regions during adulthood (Hunt et al., 2019) as observed in our study. In general, theta oscillations are believed to temporally coordinate higher-frequency activity such as gamma (Canolty et al., 2006), together supporting feedforward signaling in the visual cortex of primates (Bastos et al., 2015). Feedback communication instead relied on the beta band (Bastos et al., 2015). Thus, our connectivity findings might mirror age-related effects for visual processes in these frequency bands and coincide with an fMRI study in elderlies, which reported reduced selective responsiveness of the visual cortex to visual stimuli and slowing of perceptual speed study (Park et al., 2004). In a Cam-CAN data study, early visual areas exhibited decreased occurrence of microstates (Coquelet et al., 2020; Tibon et al., 2021). In higher-order brain areas, however, such transient neuronal states occurred more frequently with age, which correlated with lower fluid but not crystalline intelligence possibly indicating lower flexibility and coordination in the brain (Tibon et al., 2021). We also found a different pattern for age effects in higher-order regions, compatible with the notion that these areas show protracted maturation compared with lower-order regions (Sydnor et al., 2021). Notably, this was only the case for connectivity in the beta band. Here, the levels peaked around the 50s, possibly subserving intellectual, behavioral, and socioemotional processes through distributed long-range connections in the brain (Buckner and Krienen, 2013; Kopell et al., 2000; Sepulcre et al., 2010; Sydnor et al., 2021). Overall, the spatial description of MEG phase connectivity during rest across the lifespan reveals prominent effects for sensory and association regions in specific frequencies.

### 4.2. Age-related power shifts from low to high frequencies

With regards to power, we replicate a well-described shift from lower to higher frequency bands with age (Coquelet et al., 2020; Gómez et al., 2013; Hunt et al., 2019; Marek et al., 2018; Miskovic et al., 2015; Rempe et al., 2023; Whitaker et al., 2016) and report regional information on the effects. Interestingly, we found an increase with age for alpha power in temporal regions together with comparable studies using source reconstruction methods (Hunt et al., 2019; Rempe et al., 2023). Conversely, previous studies based on sensor-level data tended to indicate an age-related alpha power decrease (Polich, 1997; Thuwal et al., 2021; Tröndle et al., 2023), suggesting that the choice of analysis space is likely a source of discrepancies between studies. Overall, the linear age effects on power across frequencies were accentuated in cingulate and insular regions connected to multiple, widely different brain functions (Nieuwenhuys, 2012). These include interoception that may vary gradually with age, for example, for pain, temperature, or tactile stimuli (Jones et al., 2010), as well as temporal and social perception (Schirmer et al., 2016). Besides linear age effects, we found U-shaped trajectories in anterior-basal areas for delta, and, to some extent, also for theta power, which might be linked to the development (Campbell and Feinberg, 2009) or decline of frontal cognitive functions. For example, decline in executive functions with age is thought to be related to structural changes in the frontal lobe (Greenwood, 2000; West, 2000). Consistently, a decrease in fluid intelligence and multitasking was found in the Cam-CAN data, to which specific prefrontal changes in gray and white matter contributed (Kievit et al., 2014). Future studies should address this neurocognitive relationship across age in more detail, but the possible importance of delta oscillations for cognitive processes has been discussed previously (Harmony, 2013). Power in higher frequency bands like high beta and gamma exhibited a reverse effect with midlife peaks in cingulate and central brain regions. Cingulate regions are considered part of the DMN (Buckner et al., 2008), relate to internal cognition (Buckner et al., 2008; Raichle et al., 2001), and have been proposed to contribute to metastability of intrinsic connectivity (Leech and Sharp, 2014). Also, abnormal structure and function in this part of the cortex is associated with many neurological and psychiatric disorders with onset in adolescence and old age (Zhang and Raichle, 2010). Finally, our findings for the beta band in the central cortex might further link to altered levels of event-related activity. Central regions and beta band activity are classically related to movement (Jurkiewicz et al., 2006; Pfurtscheller and Neuper, 1997; Pfurtscheller et al., 1996) and have been shown to change across the lifespan during button pressing tasks, as measured in the Cam-CAN data set (Bardouille et al., 2019). Altogether, power alterations in our study spatially encompassed the major players of the association and sensorimotor networks with different sensitivities for age effects.

### 4.3. Divergent MEG trajectories among adult males and females

Furthermore, our results point to sex-specific effects on resting-state variability in adults in low-frequency delta and theta after accounting for intracranial volume. Our sample of 25 individuals of each sex per life decade may not yet be sufficient to detect sex effects reliably. Also, lower reliability of M/EEG estimates in the delta range and susceptibility to noise might have led to the different slopes for males and females. On the other hand, our study was age- and sex-balanced, suggesting differences in oscillations, which, as alluded to above, might play a dominant role in development and age-related decline. For example, the “occipital delta of youth” is a well-known EEG phenomenon that usually disappears with the transition to adulthood (Ebner et al., 2006) and might follow a different course between the sexes, extending into middle and late adulthood. A similar scenario might be conceivable for functional network organization, which develops from increasing integration up to early adulthood (Oldham et al., 2022; Schäfer et al., 2014) towards less within-network connectivity in old age (Damoiseaux, 2017). It should be noted though, that the interaction effects of sex in our study were limited to the low frequency range and largely mirror the overall lifespan effects found. This argues for delta connectivity differences in the trajectories putatively associated with visual processes in the occipital cortex. Variability in delta/theta power trajectories might reflect sex-related differences in cognitive frontal lobe functions as described above. Generally, similar studies in adults using MEG amplitude and phase measures have been scarce. One recent work consistently reported effects of sex in delta, theta, and alpha power across a wide age range (Rempe et al., 2023). Another study found sex-specific reconfigurations of EEG microstates during maturation and in old age, similarly pointing to different trajectories of temporal dynamics for males and females (Tomescu et al., 2018). Nonetheless, further studies should systematically focus on sex-dimorphic patterns using electrophysiology across adulthood and associate them with other phenotypes to explore behavioral associations.

### 4.4. Correspondence between MEG activity and cortical thickness across adulthood

Finally, we show that, at the cross-sectional level, the age trajectories for mainly delta and beta band oscillations spatially resembled that for cortical thickness. In general, the correspondence was of moderate effect size (*r_s_* ∼ 0.2-0.5), similar to previous attempts directly linking structural and functional connectivity using MRI (Suárez et al., 2020). Notably, variation in cortical thickness is arranged along a poster-anterior axis that is thought to have unfolded during neurogenesis and to be related to myelination (Sydnor et al., 2021) and genetic components (Valk et al., 2020). Similarly, beta band power has been expressed closely along this posterior-anterior axis, which was less stringent in the theta and alpha bands (Mahjoory et al., 2020). In addition, evidence is accumulating that cortical alterations during development also occur along a hierarchical order (Sydnor et al., 2021), as mentioned previously, for example, for delta and beta power in opposite directions in an adolescent sample (Marek et al., 2018). Thus, it is conceivable that oscillatory activity and cortical macrostructure follow spatially similar pathways during adulthood.

Moreover, cortical thickness significantly explained variance in MEG connectivity in ventral attention and default mode networks across the entire sample, hence across all age groups, and mainly in the delta to alpha frequencies. Interestingly, rapid state reconfigurations in these frequency bands have been shown to form the default mode activation pattern incorporating regions that support higher-order cognition (Vidaurre et al., 2018). The coupling between cortical thickness and low-frequency neuromagnetic connectivity might thus be the backbone of cognitive processes during adulthood. Conversely, our data revealed a negative effect of cortical thickness on high-frequency power in sensorimotor, visual, and dorsal attention networks. The cortex in sensorimotor regions is thinner than in association areas and exhibits stronger myelination that has been related to beta and gamma activation patterns before (Hunt et al., 2016), consistent with our findings. We further extend previous investigations on structure-function interrelations using MRI that have reported the opposite, that is, decoupling towards multimodal cortex but tight coupling in unimodal sensory and motor areas (Suárez et al., 2020; Vázquez-Rodríguez et al., 2019). Therefore, the relationship between brain activity at different temporal scales and macro- and microstructural features may be non-trivial. Importantly, our results may rely on mixed effects or be driven by particular age groups and should be confirmed in appropriate samples with specific age ranges. In addition, simple correlative models may not be sufficiently suited to describe complex interactions between brain structure and function. As previously suggested, microscale attributes and detailed topological information could substantially improve the modeling approaches (Suárez et al., 2020).

### 4.5. Further considerations

Our study has limitations. First, the results are based on resting-state activity with eyes closed and may be different with eyes open data, which typically are more artifactual. The eyes open scenario might lead to less alpha power in the visual cortex (Berger, 1929; Petro et al., 2022) and stronger age effects on theta power as previously reported in adolescents (Petro et al., 2022). For MEG phase-connectivity, differences between eyes open and closed conditions have been largely unexplored, but fMRI studies point to effects beyond the visual cortex (Agcaoglu et al., 2019; Patriat et al., 2013). In general, high signal power, hence a better signal-to-noise ratio, leads to a more stable phase estimation (Daffertshofer and van Wijk, 2011) and thus, potentially to a more robust estimation of age effects. This further implies non-physiological dependency between MEG power and phase-synchronization measures, but also physiological coupling has been noted (Tewarie et al., 2019). In our study, the spatial profile for the age effects on power and connectivity were mostly distinct except in the beta frequencies. Here, the age trajectories exhibited some spatial overlap and might either reflect common biological processes through coordination across distant brain regions and/or technical associations (Tewarie et al., 2019).

Moreover, what occurs as frequency-specific changes in power may be confounded by shifts in center frequencies, reductions in broadband power or changes of the aperiodic exponents (Donoghue et al., 2020). To account for this concern, we performed spectral parametrization of the broadband MEG power spectrum and overall found comparable age effects for the aperiodic-adjusted power as with the original power spectrum, pointing to true effects for oscillatory power. Please see **Suppl Figures S3** and **S4** for further details. We would like to point out that the relatively small effect sizes of the age effects on the global estimates suggest that many other factors contribute to the variance in our data. Another limitation is that the cross-sectional design of the study is not suitable for determining individual long-term trajectories (Lindenberger et al., 2011) and cohort effects cannot be excluded (Sliwinski et al., 2010).

## 5. Conclusions

Overall, we complement age-related (f)MRI literature with investigations of fast neuronal activity at rest for whole-brain networks. Source reconstruction of MEG signal characteristics yielded relevant spatial information about lifespan signatures that depend on temporal properties and partially on sex. The covariation of cortical thickness and MEG delta/beta oscillations throughout life reported here may provide an integrative imaging model for cognitive or clinical phenotypes. The associations between cortical thickness and MEG markers beyond age substantially contribute to the current perspectives on cortical structure-function relationships with reverse effects for unimodal and multimodal regions. Our findings have important implications for many studies using M/EEG. When applying similar measures in clinical cohorts, convergence or divergence from the patterns in healthy individuals could be tested, primarily to understand brain pathology and advance the development of biological disease markers.

## Data and code availability

Raw data were provided by the Cam-CAN project and are available at https://camcan-archive.mrc-cbu.cam.ac.uk/dataaccess/ under specified conditions. Main processed MEG and structural data and all code used for the pre-processing, statistical analyses, and visualization of the results have been deposited in https://github.com/chstier/Lifespan_MEG.

## Credit authorship contribution statement

**Christina Stier:** Conceptualization, Methodology, Formal analysis, Writing – original draft, Writing – review & editing, Visualization. **Christoph Braun:** Methodology, Writing – review & editing. **Niels K. Focke:** Conceptualization, Methodology, Writing – review & editing, Supervision

## Acknowledgments

Data collection and sharing for this project was provided by the Cambridge Centre for Ageing and Neuroscience (CamCAN). Cam-CAN funding was provided by the UK Biotechnology and Biological Sciences Research Council (grant number BB/H008217/1), together with support from the UK Medical Research Council and University of Cambridge, UK. This work was further supported by the German Research Foundation (DFG; grant number FO 750/5-1 to N.K.F.)

## Supporting Information

**Fig. S1.**
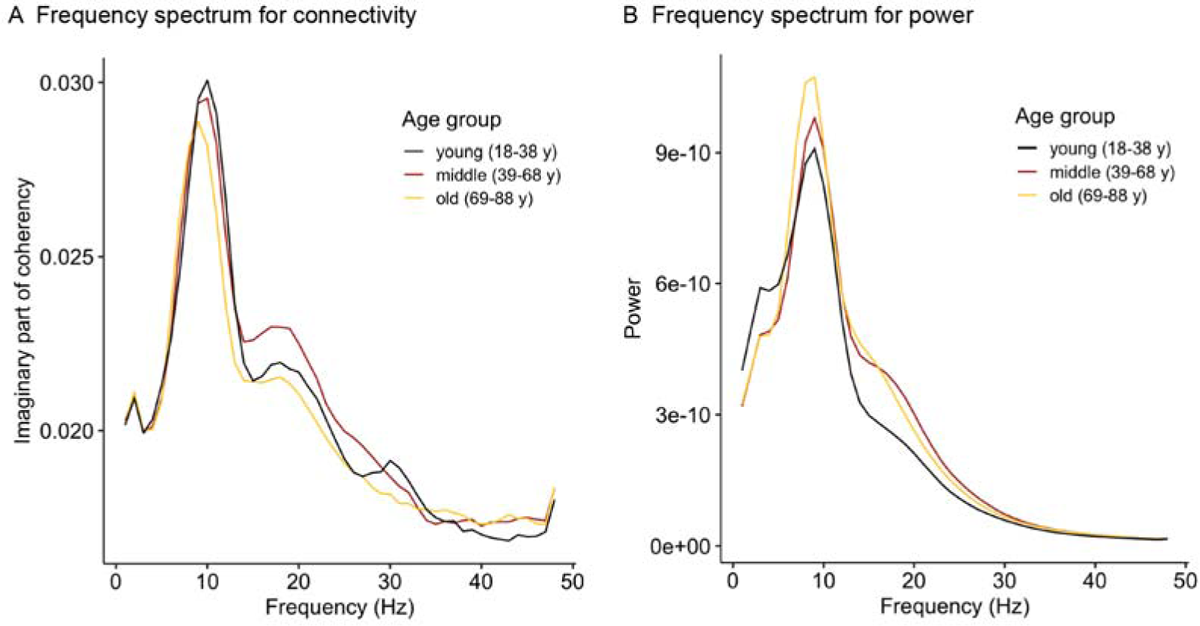
Spectra for connectivity and power across age groups For each individual included in this study, global connectivity or power (fT^2^) was computed for 1 Hz frequency bins and averaged for the corresponding age group (*n*_young_ = 100, *n*_middle_ = 150, *n*_old_ = 100). Black colors represent the young age group, red represents the middle age group, and light colors indicate the old age group.

**Fig. S2.**
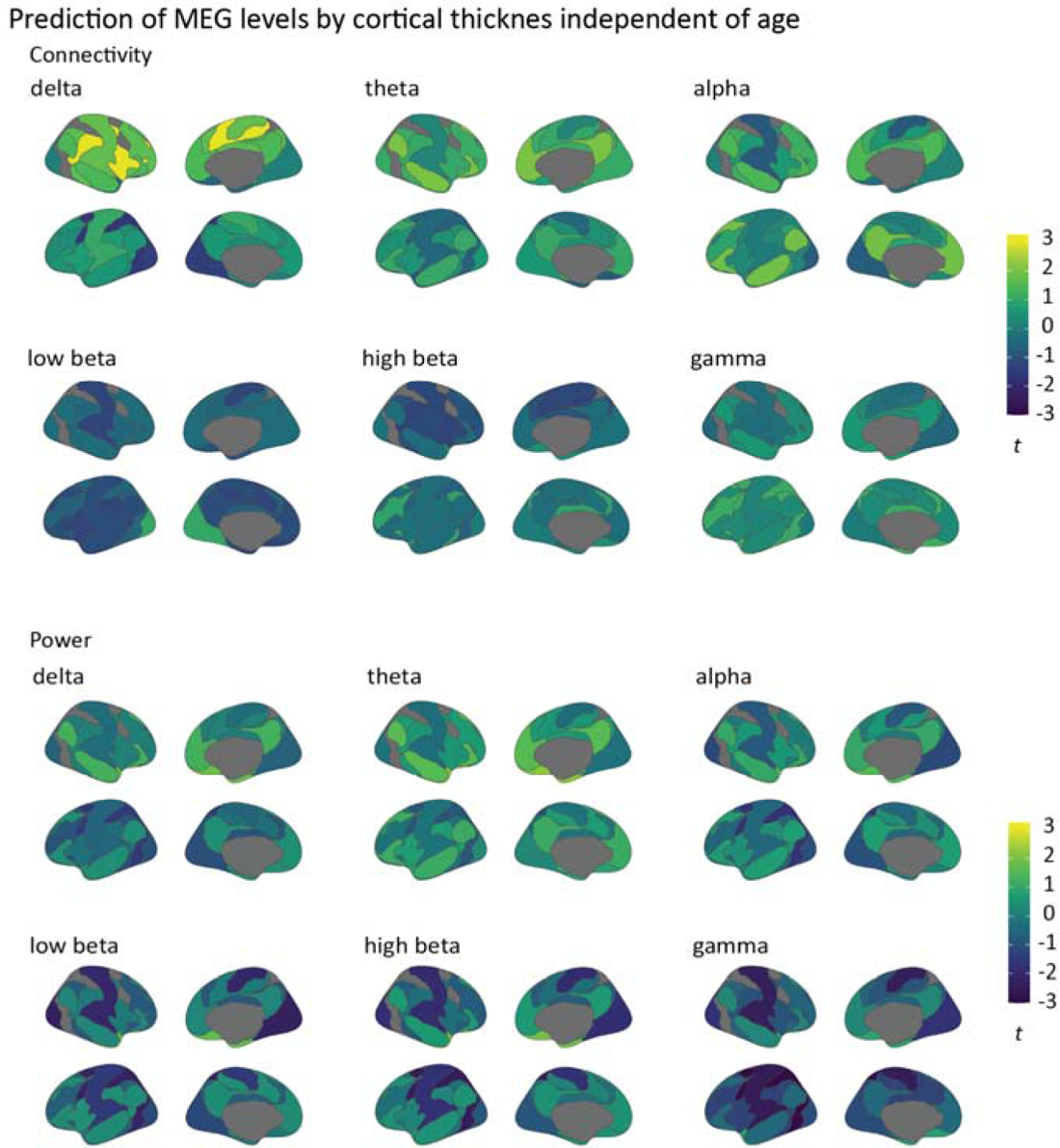
Full *t*-value maps for the effect of cortical thickness on MEG levels across the adult lifespan sample Shown is the influence of cortical thickness on functional MEG markers (t-values) across the whole study sample after regressing out the effects of covariables. We estimated general linear models including cortical thickness, age, age^2^, sex, and total intracranial volume as independent variables and employed a statistical permutation framework to the models at the level of functional resting-state networks, as described in Yeo et al. (2011). Please see the main manuscript for the results after significance testing.

**Fig. S3.**
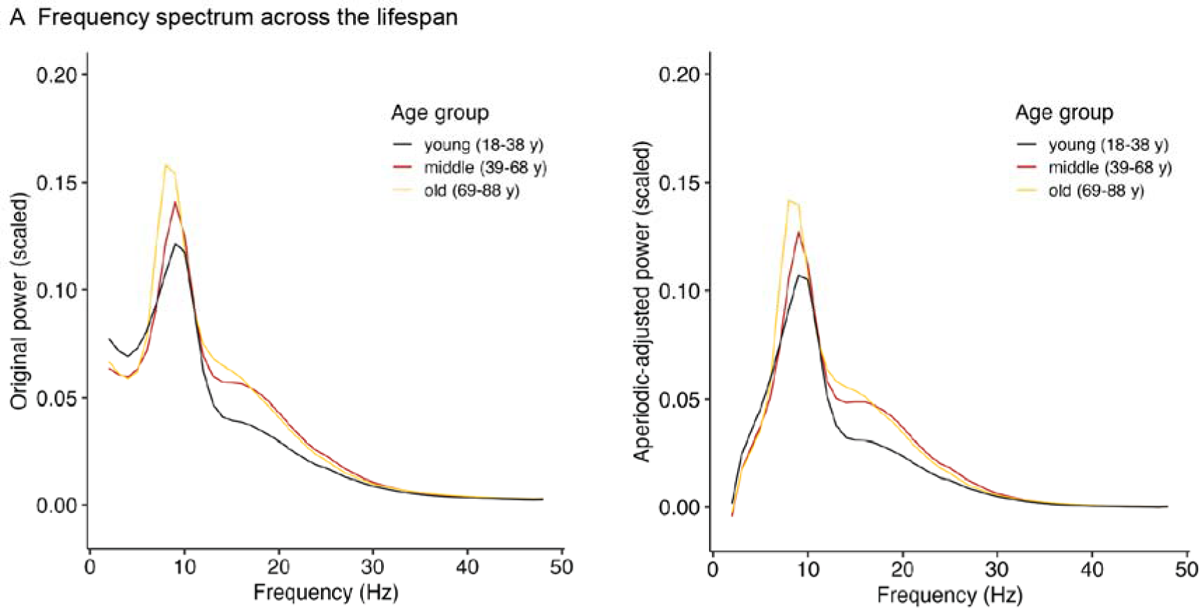
Secondary analyses: original power and aperiodic-adjusted power spectra We used the specparam algorithm (Donoghue et al., 2020) to parameterize the power spectrum at the source-level after applying unit-noise gain linear constrained minimum variance (LCMV) beamforming (Van Veen et al., 1997). Using LCMV spatial filters were constructed (regularization lambda = 5%, free-orientation forward solution) and multiplied with the sensor time series data, resulting in source level time series. To keep the computational burden low, we used 200 parcels of the Schäfer atlas (Schaefer et al., 2018) as source model instead of the high-resolution mesh with 2004 vertices as used throughout the main analyses. We then passed the frequency range of 1-60 Hz of the source power spectrum to the specparam algorithm with the following settings: peak width limits [1-8]; maximum number of peaks: infinitive; minimum peak height: 0; and aperiodic mode: fixed. This resulted in aperiodic and periodic (“oscillatory”) signal components for each subject. The plot shows the original and aperiodic-adjusted frequency spectrum at each frequency bin averaged for the corresponding age group (*n*_young_ = 100, *n*_middle_ = 150, *n*_old_ = 100). Black colors represent the young age group, red represents the middle age group, and light colors indicate the old age group. Note that power values (fT^2^) were scaled with the factor 1*e^13^ before subjected to the beamformer.

**Fig. S4.**
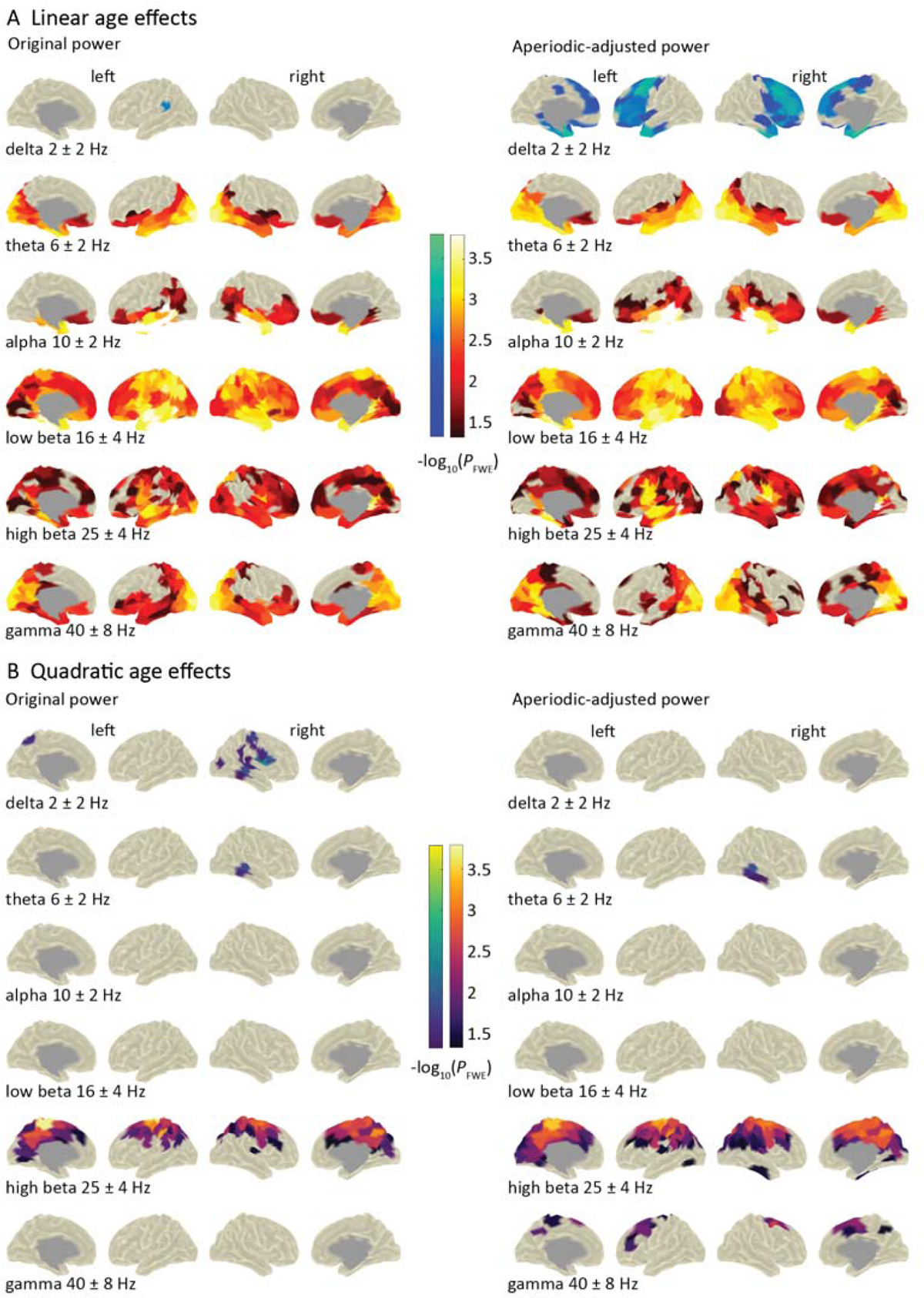
Secondary analyses: age effects on original and aperiodic-adjusted power To evaluate whether the estimation of age effects on the original and the aperiodic-adjusted power differ, we used permutation analysis of linear models at the level of the Schäfer parcels (Schaefer et al., 2018). Individual power values were averaged for the respective frequency ranges. Analogue to the primary analyses of this work, we considered age, age^2^, sex, and total intracranial volume as independent variables. The significance level was set at -log10 *p* > 1.3 (equivalent to *p* < 0.05), family-wise error corrected (FWE). Left = left hemisphere, right = right hemisphere. (A) The red color bar indicates significant positive effects of age and blue indicates negative effects of age. (B) The green color bar depicts significant quadratic effects of age following a U-shaped pattern (convex), while the purple color bar indicates significant effects following an inverted U (concave). We would like to add that the fitting using the specparam algorithm for very low frequencies can be biased and should be investigated further. Please note that in general, minor differences between the results for the original power in the main analysis and the one presented here can be expected due to the use of different beamforming methods and spatial resolution (vertex-versus atlas-based). Specifically, the spatial filters using DICS beamforming methods as in our main analysis are optimized for specific frequency ranges whereas the LCMV beamformer was built on the broadband signal.

---

**Table S1.** Demographic data of the individual age groups included in the study

| Age group | *N* | Age (years) | Sex |
| --- | --- | --- | --- |
| | | Mean ± SD | Males : females |
| 18-28 years | 50 | 24.4 ± 3.4 | 25 : 25 |
| 29-38 years | 50 | 33.3 ± 2.9 | 25 : 25 |
| 39-48 years | 50 | 43.6 ± 3.0 | 25 : 25 |
| 49-58 years | 50 | 53.7 ± 2.8 | 25 : 25 |
| 59-68 years | 50 | 63.0 ± 2.8 | 25 : 25 |
| 69-78 years | 50 | 71.8 ± 2.9 | 25 : 25 |
| 79-88 years | 50 | 81.5 ± 2.7 | 25 : 25 |

**Table S2.**
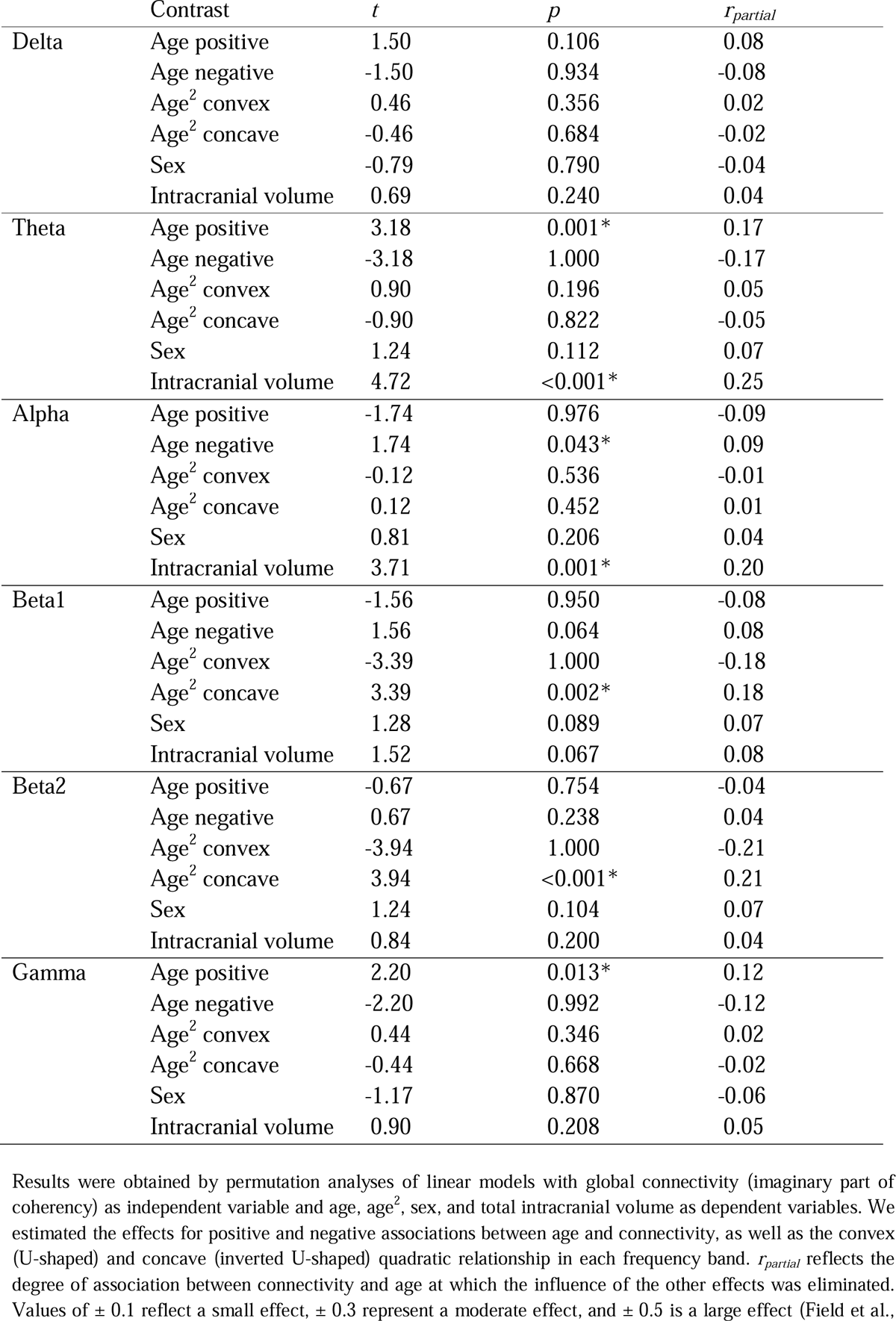
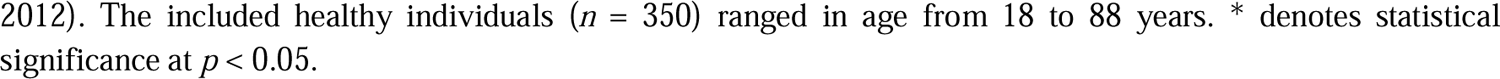
Relationship between global connectivity and age

|  | Contrast | *t* | *p* | *r*<sub>partial</sub> |
| --- | --- | --- | --- | --- |
| Delta | Age positive | 1.50 | 0.106 | 0.08 |
|  | Age negative | -1.50 | 0.934 | -0.08 |
|  | Age<sup>2</sup> convex | 0.46 | 0.356 | 0.02 |
|  | Age<sup>2</sup> concave | -0.46 | 0.684 | -0.02 |
|  | Sex | -0.79 | 0.790 | -0.04 |
|  | Intracranial volume | 0.69 | 0.240 | 0.04 |
| Theta | Age positive | 3.18 | 0.001* | 0.17 |
|  | Age negative | -3.18 | 1.000 | -0.17 |
|  | Age<sup>2</sup> convex | 0.90 | 0.196 | 0.05 |
|  | Age<sup>2</sup> concave | -0.90 | 0.822 | -0.05 |
|  | Sex | 1.24 | 0.112 | 0.07 |
|  | Intracranial volume | 4.72 | <0.001* | 0.25 |
| Alpha | Age positive | -1.74 | 0.976 | -0.09 |
|  | Age negative | 1.74 | 0.043* | 0.09 |
|  | Age<sup>2</sup> convex | -0.12 | 0.536 | -0.01 |
|  | Age<sup>2</sup> concave | 0.12 | 0.452 | 0.01 |
|  | Sex | 0.81 | 0.206 | 0.04 |
|  | Intracranial volume | 3.71 | 0.001* | 0.20 |
| Beta1 | Age positive | -1.56 | 0.950 | -0.08 |
|  | Age negative | 1.56 | 0.064 | 0.08 |
|  | Age<sup>2</sup> convex | -3.39 | 1.000 | -0.18 |
|  | Age<sup>2</sup> concave | 3.39 | 0.002* | 0.18 |
|  | Sex | 1.28 | 0.089 | 0.07 |
|  | Intracranial volume | 1.52 | 0.067 | 0.08 |
| Beta2 | Age positive | -0.67 | 0.754 | -0.04 |
|  | Age negative | 0.67 | 0.238 | 0.04 |
|  | Age<sup>2</sup> convex | -3.94 | 1.000 | -0.21 |
|  | Age<sup>2</sup> concave | 3.94 | <0.001* | 0.21 |
|  | Sex | 1.24 | 0.104 | 0.07 |
|  | Intracranial volume | 0.84 | 0.200 | 0.04 |
| Gamma | Age positive | 2.20 | 0.013* | 0.12 |
|  | Age negative | -2.20 | 0.992 | -0.12 |
|  | Age<sup>2</sup> convex | 0.44 | 0.346 | 0.02 |
|  | Age<sup>2</sup> concave | -0.44 | 0.668 | -0.02 |
|  | Sex | -1.17 | 0.870 | -0.06 |
|  | Intracranial volume | 0.90 | 0.208 | 0.05 |

Results were obtained by permutation analyses of linear models with global connectivity (imaginary part of coherency) as independent variable and age, age<sup>2</sup>, sex, and total intracranial volume as dependent variables. We estimated the effects for positive and negative associations between age and connectivity, as well as the convex (U-shaped) and concave (inverted U-shaped) quadratic relationship in each frequency band. *r*<sub>partial</sub> reflects the degree of association between connectivity and age at which the influence of the other effects was eliminated. Values of $\pm 0.1$ reflect a small effect, $\pm 0.3$ represent a moderate effect, and $\pm 0.5$ is a large effect (Field et al.,
2012). The included healthy individuals ( $n = 350$ ) ranged in age from 18 to 88 years. \* denotes statistical significance at $p < 0.05$ .

**Table S3.**
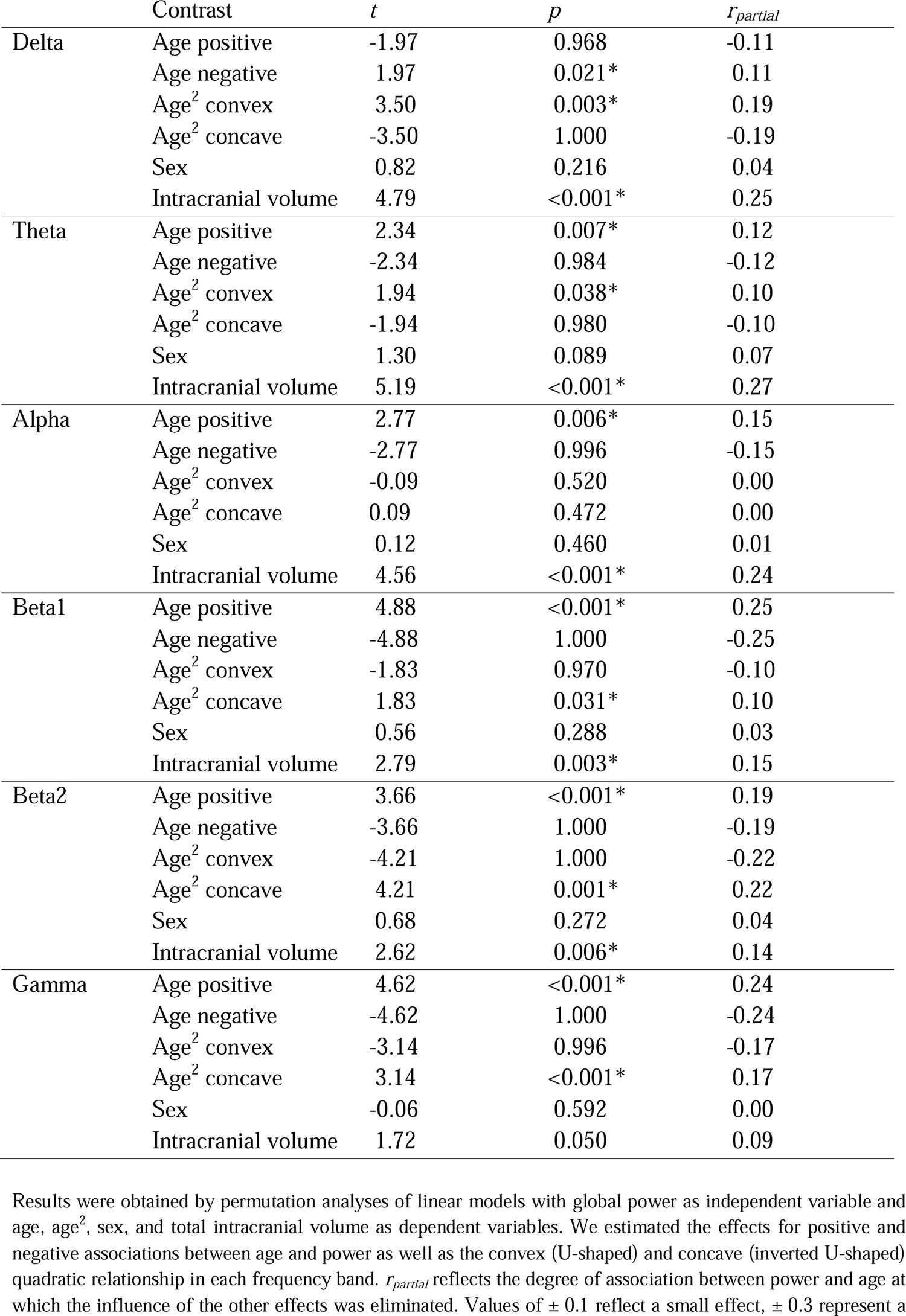
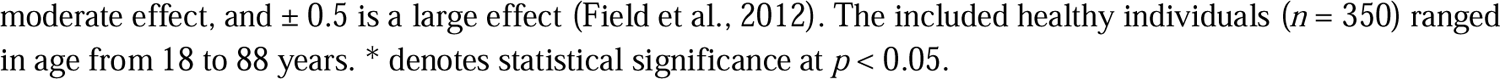
Relationship between global power and age

|  | Contrast | *t* | *p* | *r<sub>partial</sub>* |
| --- | --- | --- | --- | --- |
| Delta | Age positive | -1.97 | 0.968 | -0.11 |
|  | Age negative | 1.97 | 0.021* | 0.11 |
|  | Age<sup>2</sup> convex | 3.50 | 0.003* | 0.19 |
|  | Age<sup>2</sup> concave | -3.50 | 1.000 | -0.19 |
|  | Sex | 0.82 | 0.216 | 0.04 |
|  | Intracranial volume | 4.79 | <0.001* | 0.25 |
| Theta | Age positive | 2.34 | 0.007* | 0.12 |
|  | Age negative | -2.34 | 0.984 | -0.12 |
|  | Age<sup>2</sup> convex | 1.94 | 0.038* | 0.10 |
|  | Age<sup>2</sup> concave | -1.94 | 0.980 | -0.10 |
|  | Sex | 1.30 | 0.089 | 0.07 |
|  | Intracranial volume | 5.19 | <0.001* | 0.27 |
| Alpha | Age positive | 2.77 | 0.006* | 0.15 |
|  | Age negative | -2.77 | 0.996 | -0.15 |
|  | Age<sup>2</sup> convex | -0.09 | 0.520 | 0.00 |
|  | Age<sup>2</sup> concave | 0.09 | 0.472 | 0.00 |
|  | Sex | 0.12 | 0.460 | 0.01 |
|  | Intracranial volume | 4.56 | <0.001* | 0.24 |
| Beta1 | Age positive | 4.88 | <0.001* | 0.25 |
|  | Age negative | -4.88 | 1.000 | -0.25 |
|  | Age<sup>2</sup> convex | -1.83 | 0.970 | -0.10 |
|  | Age<sup>2</sup> concave | 1.83 | 0.031* | 0.10 |
|  | Sex | 0.56 | 0.288 | 0.03 |
|  | Intracranial volume | 2.79 | 0.003* | 0.15 |
| Beta2 | Age positive | 3.66 | <0.001* | 0.19 |
|  | Age negative | -3.66 | 1.000 | -0.19 |
|  | Age<sup>2</sup> convex | -4.21 | 1.000 | -0.22 |
|  | Age<sup>2</sup> concave | 4.21 | 0.001* | 0.22 |
|  | Sex | 0.68 | 0.272 | 0.04 |
|  | Intracranial volume | 2.62 | 0.006* | 0.14 |
| Gamma | Age positive | 4.62 | <0.001* | 0.24 |
|  | Age negative | -4.62 | 1.000 | -0.24 |
|  | Age<sup>2</sup> convex | -3.14 | 0.996 | -0.17 |
|  | Age<sup>2</sup> concave | 3.14 | <0.001* | 0.17 |
|  | Sex | -0.06 | 0.592 | 0.00 |
|  | Intracranial volume | 1.72 | 0.050 | 0.09 |

Results were obtained by permutation analyses of linear models with global power as independent variable and age, age<sup>2</sup>, sex, and total intracranial volume as dependent variables. We estimated the effects for positive and negative associations between age and power as well as the convex (U-shaped) and concave (inverted U-shaped) quadratic relationship in each frequency band. *r<sub>partial</sub>* reflects the degree of association between power and age at which the influence of the other effects was eliminated. Values of $\pm 0.1$ reflect a small effect, $\pm 0.3$ represent a
moderate effect, and $\pm 0.5$ is a large effect (Field et al., 2012). The included healthy individuals ( $n = 350$ ) ranged in age from 18 to 88 years. \* denotes statistical significance at $p < 0.05$ .

**Table S4.** Summary of interaction effects sex by age on global MEG metrics

|  |  | Connectivity |  | Power |  |
| --- | --- | --- | --- | --- | --- |
|  | Contrast | *t* | *p* | *t* | *p* |
| Delta | Sex*age positive | -1.24 | 0.882 | -2.77 | 0.996 |
|  | Sex*age negative | 1.24 | 0.108 | 2.77 | 0.007* |
|  | Sex*age<sup>2</sup> convex | -2.06 | 0.988 | 0.42 | 0.318 |
|  | Sex*age<sup>2</sup> concave | 2.06 | 0.017* | -0.42 | 0.672 |
| Theta | Sex*age positive | 0.76 | 0.204 | -1.54 | 0.928 |
|  | Sex*age negative | -0.76 | 0.762 | 1.54 | 0.064 |
|  | Sex*age<sup>2</sup> convex | 0.55 | 0.286 | 0.52 | 0.280 |
|  | Sex*age<sup>2</sup> concave | -0.55 | 0.686 | -0.52 | 0.698 |
| Alpha | Sex*age positive | 0.85 | 0.200 | -0.76 | 0.752 |
|  | Sex*age negative | -0.85 | 0.772 | 0.76 | 0.234 |
|  | Sex*age<sup>2</sup> convex | -1.49 | 0.932 | 0.13 | 0.448 |
|  | Sex*age<sup>2</sup> concave | 1.49 | 0.062 | -0.13 | 0.578 |
| Beta1 | Sex*age positive | 0.74 | 0.240 | -0.71 | 0.768 |
|  | Sex*age negative | -0.74 | 0.772 | 0.71 | 0.246 |
|  | Sex*age<sup>2</sup> convex | 0.32 | 0.362 | 0.04 | 0.458 |
|  | Sex*age<sup>2</sup> concave | -0.32 | 0.614 | -0.04 | 0.558 |
| Beta2 | Sex*age positive | -1.14 | 0.908 | -0.73 | 0.784 |
|  | Sex*age negative | 1.14 | 0.130 | 0.73 | 0.240 |
|  | Sex*age<sup>2</sup> convex | -0.14 | 0.564 | -0.02 | 0.504 |
|  | Sex*age<sup>2</sup> concave | 0.14 | 0.390 | 0.02 | 0.526 |
| Gamma | Sex*age positive | -0.55 | 0.694 | -0.19 | 0.544 |
|  | Sex*age negative | 0.55 | 0.300 | 0.19 | 0.418 |
|  | Sex*age<sup>2</sup> convex | 1.43 | 0.068 | -0.14 | 0.534 |
|  | Sex*age<sup>2</sup> concave | -1.43 | 0.916 | 0.14 | 0.420 |

Results were obtained by permutation analyses of linear models testing whether the coefficients for the age and age<sup>2</sup> effect on MEG metrics differed between males ( $n = 175$ ) and females ( $n = 175$ ) after correcting for the influence of total intracranial volume. Positive and negative linear associations between age and the metrics as well as the convex (U-shaped) and concave (inverted U-shaped) quadratic relationship in each frequency band were considered. The included healthy individuals ( $n = 350$ ) ranged in age from 18 to 88 years. \* denotes statistical significance at $p < 0.05$ .

**Table S5.** Correlation between (negative) linear age effects on cortical thickness and MEG markers

| | Metric | Age effect | $r_s$ | $p_{spin}$ |
| --- | --- | --- | --- | --- |
| Delta | Connectivity | Linear (positive) | .343 | 0.004* |
|  | Connectivity | Quadratic (convex) | .100 | 0.561 |
|  | Power | Linear (negative) | .314 | 0.008* |
|  | Power | Quadratic (convex) | - .014 | 0.961 |
| Theta | Connectivity | Linear (positive) | .030 | 0.932 |
|  | Connectivity | Quadratic (concave) | .205 | 0.351 |
|  | Power | Linear (positive) | - .028 | 0.935 |
|  | Power | Quadratic (convex) | - .152 | 0.568 |
| Alpha | Connectivity | Linear (negative) | .012 | 0.967 |
|  | Connectivity | Quadratic (convex) | .220 | 0.095 |
|  | Power | Linear (positive) | .183 | 0.499 |
|  | Power | Quadratic (convex) | - .243 | 0.249 |
| Beta1 | Connectivity | Linear (negative) | - .015 | 0.963 |
|  | Connectivity | Quadratic (concave) | .227 | 0.197 |
|  | Power | Linear (positive) | .303 | 0.104 |
|  | Power | Quadratic (concave) | .431 | 0.007* |
| Beta2 | Connectivity | Linear (negative) | - .181 | 0.581 |
|  | Connectivity | Quadratic (concave) | .267 | 0.253 |
|  | Power | Linear (positive) | .275 | 0.139 |
|  | Power | Quadratic (concave) | .479 | 0.002* |
| Gamma | Connectivity | Linear (positive) | - .051 | 0.851 |
|  | Connectivity | Quadratic (convex) | - .052 | 0.781 |
|  | Power | Linear (positive) | .093 | 0.495 |
|  | Power | Quadratic (convex) | - .366 | 0.077 |

Correlation analyses were performed between the age effect $t$ statistics on cortical thickness and those on MEG markers at the vertex-level. Statistical significance of the correlation coefficients was tested using 10,000 random rotations (spin test; Alexander-Bloch et al. (2018)). Significant correlations ( $p_{spin} < 0.05$ ) suggest spatial correspondence of MEG alterations with cortical thinning across the lifespan ( $r_s$ = Spearman rank correlation coefficient; $r_s < .3$ indicates a weak, $r_s$ from .3 to .5 indicates a moderate relationship (Field et al., 2012)).

**Table S6.** Correlation between quadratic age effects (concave) on cortical thickness and MEG markers

| | Metric | Age effect | $r_s$ | $p_{spin}$ |
| --- | --- | --- | --- | --- |
| Delta | Connectivity | Linear (positive) | .053 | 0.719 |
|  | Connectivity | Quadratic (convex) | - .218 | 0.028* |
|  | Power | Linear (negative) | - .016 | 0.855 |
|  | Power | Quadratic (convex) | .113 | 0.543 |
| Theta | Connectivity | Linear (positive) | .298 | 0.086 |
|  | Connectivity | Quadratic (concave) | - .049 | 0.771 |
|  | Power | Linear (positive) | .289 | 0.109 |
|  | Power | Quadratic (convex) | .147 | 0.441 |
| Alpha | Connectivity | Linear (negative) | .295 | 0.050 |
|  | Connectivity | Quadratic (convex) | - .109 | 0.313 |
|  | Power | Linear (positive) | .121 | 0.553 |
|  | Power | Quadratic (convex) | .107 | 0.548 |
| Beta1 | Connectivity | Linear (negative) | .360 | 0.024* |
|  | Connectivity | Quadratic (concave) | .193 | 0.118 |
|  | Power | Linear (positive) | .005 | 0.975 |
|  | Power | Quadratic (concave) | - .065 | 0.727 |
| Beta2 | Connectivity | Linear (negative) | .218 | 0.386 |
|  | Connectivity | Quadratic (concave) | - .144 | 0.441 |
|  | Power | Linear (positive) | - .012 | 0.936 |
|  | Power | Quadratic (concave) | - .174 | 0.321 |
| Gamma | Connectivity | Linear (positive) | .027 | 0.897 |
|  | Connectivity | Quadratic (convex) | <.001 | 0.996 |
|  | Power | Linear (positive) | .038 | 0.671 |
|  | Power | Quadratic (convex) | .171 | 0.351 |

Correlation analyses were performed between the age effect $t$ statistics on cortical thickness and those on MEG markers at the vertex-level. Statistical significance of the correlation coefficients was tested using 10,000 random rotations (spin test; Alexander-Bloch et al. (2018)). Significant correlations ( $p_{spin} < 0.05$ ) suggest spatial correspondence of MEG and cortical thickness alterations across the lifespan ( $r_s$ = Spearman rank correlation coefficient; $r_s < .3$ indicates a weak, $r_s$ from .3 to .5 indicates a moderate relationship (Field et al., 2012)).

